# Spatial community variability: Interactive effects of predators and isolation on stochastic community assembly

**DOI:** 10.1101/2020.12.22.423949

**Authors:** Rodolfo Mei Pelinson, Mathew A. Leibold, Luis Schiesari

## Abstract

In the absence of environmental heterogeneity, spatial variation among local communities can be mostly attributed to demographic stochasticity (*i*.*e*., ecological drift) and historical contingency in colonization (*i*.*e*. random dispersal and priority effects). The consequences of demographic stochasticity are highly dependent on community size, gamma, and alpha diversity, which, along with historical contingency, can be strongly affected by dispersal limitation and the presence of predators. We used freshwater insect communities to experimentally test whether and how the presence of a generalist predatory fish and dispersal limitation (*i*.*e*., isolation by distance from a source habitat) can change the relative importance of stochastic and non-stochastic processes on community variability. We found that dispersal limitation can have both negative and positive effects on community variability, and their importance may depend on the presence of predatory fish. Negative effects happened because predatory insects cannot always successfully colonize highly isolated ponds, causing herbivores and detritivores to increase in abundance. As a consequence, community size increases, decreasing the importance of demographic stochasticity on community structure. However, when fish is absent, these effects are counterbalanced by an increase in the importance of priority effects generating more distinct communities in more isolated ponds. Such effects can be caused by either pre or post-colonization mechanisms and were absent in the presence of fish predators, likely because fish prevented both predatory and non-predatory insects from becoming more abundant, irrespective of their order of colonization.

## INTRODUCTION

Ecologists have traditionally considered that environmental variation in local conditions is the main driver of differences among local community structures within metacommunities (Vellend 2016). Yet, considerable variation in community composition and abundance patterns arises even among communities found under similar environmental conditions. Such variability has been either attributed to demographic stochasticity (*i*.*e*., stochastic events of birth and death; ecological drift) (Connor and Simberloff 1979, Hubbell 2001, Vellend 2016), or historical contingency in colonization, which have both stochastic and deterministic components (Fukami 2015). The stochastic one is attributed to the degree of randomness in dispersal and colonization sequence (Vellend et al. 2014, Fukami 2015). The deterministic one is due to species interactions, which can lead communities to different deterministic structures, depending on colonization sequence (*e*.*g*., priority effects) (Diamond 1975, Law and Morton 1993, Fukami 2015). Therefore, assuming that communities within a metacommunity have similar contemporary and past abiotic conditions, the importance of deterministic and stochastic processes on community variability may vary depending on different community and metacommunity aspects, such as the size of the species pool (*i*.*e*. gamma-diveristy), local richness (*i*.*e*. alpha-diveristy), community size (*i*.*e*. total number of individuals), stochasticity in dispersal, and strength of species interactions (Chase 2003, Vellend et al. 2014, Vellend 2016, Leibold and Chase 2018).

Community variability is often referred to as beta-diversity (Anderson et al. 2011), which can be measured in different ways either emphasizing differences in species abundance patterns, or differences in species composition, giving more or less weight to rare species (Koleff et al. 2003, Barwell et al. 2015). In all cases, beta-diversity connects gamma (*i*.*e*., regional metacommunity diversity) to alfa-diversity (*i*.*e*., local community diversity) (Whittaker 1960, 1972, Anderson et al. 2011). A direct consequence of this connection is that community variability can be enhanced or dampened by any processes affecting gamma or alpha diversity (Mouquet and Loreau 2003, Chase 2007, 2010, Chase et al. 2009). For instance, processes that increase regional but not local richness can result in higher variability simply because they increase the set of possible different community compositions (*i*.*e*., statistical inevitability) (Chase et al. 2009, Chase and Myers 2011). Community size (*i*.*e*., the total number of individuals in a community) is also important in this context (Orrock and Fletcher Jr. 2005). Small populations are more likely affected by stochastic events of birth and death than larger populations. Thus, metacommunities where species are less abundant tend to have higher community variability (Myers et al. 2015, Siqueira et al. 2020).

If species interactions are important in a given biological system, the order of arrival of species should have a strong deterministic effect on community structure because the species that arrive first at a given habitat has an advantage over other competitors, apparent competitors, or even predators (Chase 2003, Shurin et al. 2004, Fukami 2015). In this case, if the identity of the first colonizers of a community is sufficiently random, the colonization sequence in different communities will follow different deterministic orders and communities will differ more than would be expected by simple demographic stochasticity or random dispersal history (Vellend et al. 2014, Fukami 2015). However, if the order of the first colonizers is deterministic (*i*.*e*. same order in multiple communities), their structures will be more similar to each other than would be expected by demographic stochasticity or random dispersal history (Vellend et al. 2014).

The consequences of stochastic processes and deterministic historical contingency on community variability can be changed by the presence of top predators in many different ways, depending on the intensity of predation and how generalist these predators are (reviewed in Appendix S1). In freshwater ponds, fish tend to be the top predators of the community (Wellborn et al. 1996). Because they are usually visually oriented predators, they preferentially prey upon more conspicuous taxa which frequently are large-bodied predatory insects (Diehl 1992, Goyke and Hershey 1992, Wellborn et al. 1996, Batzer et al. 2000, Pelinson et al. 2020). Therefore, fish can exclude more vulnerable species from the metacommunity, decreasing gamma, and consequently, beta-diversity (Chase et al. 2009). However, in the absence of its preferred prey, fish might also prey upon whatever taxa are most abundant, including herbivores and detritivores (Pelinson et al. 2020). Thus, if species interactions are important and the most abundant taxa (*i*.*e*. competitive dominants) are those who colonize a pond first, there will be a trade-off between competitive ability (determined by order of arrival) and vulnerability to predation (abundance) (Leibold 1996, 1999, Louette and De Meester 2007). In this scenario, fish may also decrease the importance of colonization sequence on community structure by suppressing competitive dominants.

Dispersal limitation can also have different non-mutually exclusive consequences to the importance of stochastic processes and deterministic historical contingency on community variability, depending on how much species vary in dispersal rates (reviewed in Appendix S1). Freshwater insects are known to vary in their ability to colonize differently isolated habitats (Shulman and Chase 2007, Chase and Shulman 2009, Hein and Gillooly 2011, Pelinson et al. 2020). Predatory insects generally have smaller population sizes and greater generation times if compared to their prey, thus they have fewer events of dispersal than herbivores and detritivores (Shulman and Chase 2007, Chase and Shulman 2009, Hein and Gillooly 2011, Pelinson et al. 2020). The consequence is that some of those predatory insects may be absent from highly isolated ponds, decreasing gamma (Hendrickx et al. 2009) and consequently, beta-diversity (Chase and Myers 2011). Additionally, if species interactions are strong, unequal dispersal rates can make beta-diversity even lower because the order of arrival of species in all communities would be determined by dispersal ability or frequency (Vellend et al. 2014).

One way to tease apart the effect of deterministic processes affecting community variability from stochastic ones is by using null models to compute how much community variability would be expected in randomly reassembled communities under the same community size, gamma, and alpha diversity (Chase et al. 2011). In this case, the deviations of the observed beta-diversity from the expected, (a metric that is termed beta-deviation), would be caused by any process not included in the null model, such as priority effects or environmental stochasticity (Chase et al. 2011, Kraft et al. 2011, Vannette and Fukami 2017).

Here we experimentally tested how the presence of a predatory fish and dispersal limitation can change the relative importance of stochastic (*e*.*g*. ecological drift and stochastic dispersal) and non-stochastic processes (*e*.*g*., priority effects) on community variability. (H1) We hypothesize that fish can decrease community variability by preferentially preying on a defined subset of prey, thus decreasing gamma diversity, and promoting the homogenization of communities (Fig. 1a and 1c). (H2) We also expect highly isolated habitats to have lower community variability because poor dispersers would not be able to reach such patches frequently (Fig. 1b and 1c). (H3) If colonization sequence and species interactions are important in determining community structure, spatial isolation should make community variability lower than would be expected by simple demographic stochasticity or random dispersal because the order of arrival of species will be determined by dispersal rates (Fig. 1d and 1f). (H4) Finally, even though fish preferentially prey upon large-bodied predatory insects, they might also substantially prey upon whatever taxa are most abundant (Pelinson et al. 2020), thus if species interactions are important, the presence of fish should decrease their importance by reducing the abundance of any taxa who successfully colonize ponds first (Fig. 1e and 1f).

**Figure 1.**
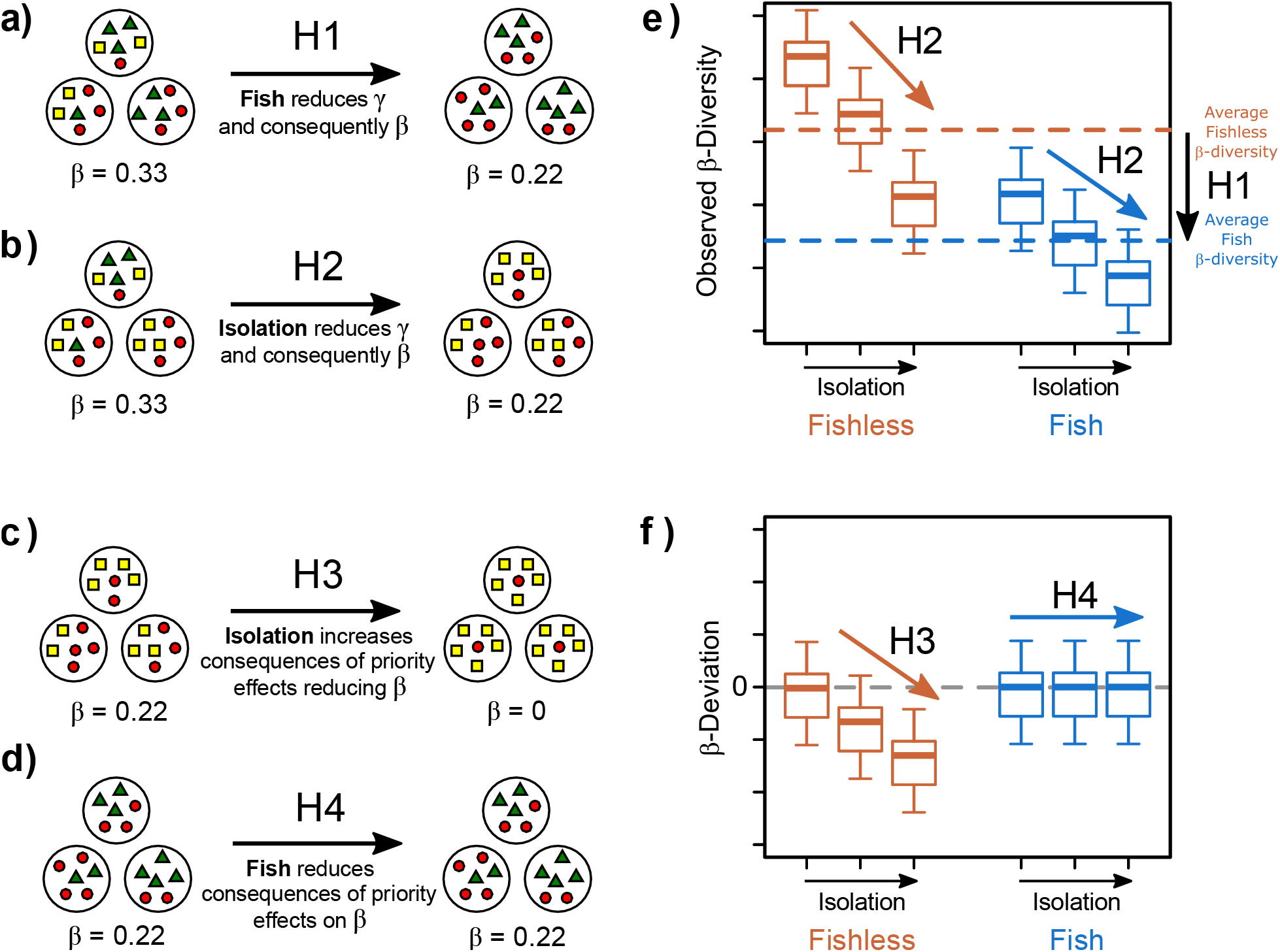
Hypotheses of how the presence of fish and spatial isolation can affect freshwater insect community variability under the same environmental conditions. (a): The presence of fish prevents successes in colonization of more vulnerable prey, reducing gamma diversity. Consequently, beta-diversity decreases as a statistical inevitability (H1). (b): Similarly, spatial isolation prevents worst dispersers from colozing isolated ponds, reducing gamma and beta-diversity (H2). (c): Spatial isolation increases the differences in frequency of arrival of bad and good dispersers, increasing the consequences of priority effects, reducing community variability (H3). (d): The presence of fish reduces the consequences of priority effects caused by isolation by preying upon the most abundant insects (H4). Right panels: *A priori* expectations of observed (e) beta-diversity and (f) beta-deviation (i.e. how much observed beta-diversity deviates from stochastic assembly) for each isolation distances in ponds with (blue boxes) and without fish (orange boxes).

## METHODS

We conducted a field experiment at the Estação Ecológica de Santa Bárbara (EESB) in Águas de Santa Bárbara, São Paulo, Brazil (22°48’59” S, 49°14’12” W). The EESB is a 2,712-ha protected area predominantly covered with open savanna vegetation, with smaller portions of seasonal semideciduous forests, *Pinus* sp., and *Eucalyptus* sp plantations (Melo and Durigan 2011). Soils are sandy and the climate is Koeppen’s Cwa, i.e., warm temperate with dry winters and hot summers (Kottek et al. 2006). Mean annual rainfall is ∼1250mm with a distinct rainy season from October to March (January being the wettest month with ∼200mm rainfall) and a dry season from April to September (July being the driest month with ∼25mm rainfall; (Hijmans et al. 2005). In the EESB the experiment was implemented in an area covered by second-growth Cerrado *sensu stricto*, a moderately dense, open-canopy savanna habitat (Melo and Durigan 2011).

Experimental units consisted of ∼1,200L artificial ponds dug into the ground and lined with a 0.5 mm thick, high-density polyethylene geomembrane to retain water. Each pond was 4m long, 1m wide, and 40 cm deep. Walls were vertical along the pond; 1m-long ramps terminating at ground level at each short side of the pond provided shallow microhabitats for freshwater organisms and escape for terrestrial fauna that eventually fell into the water. Two roof tiles were placed at the waterline in each of the short sides to provide shelter and oviposition habitat for insects. Three 30 cm-long, ten cm-wide PVC pipes were placed in the water to provide shelter for fishes. The experiment followed a fully factorial design crossing fish presence (presence/absence) with spatial isolation (three levels of isolation). The manipulated fish was the Redbreast Tilapia (*Coptodon rendalli*, standard length 99.2 mm ± 5.9 mm, wet mass 40.2 g ± 8.8 g, mean ± SD) at a density of one individual per pond. The isolation treatment was achieved by establishing eight artificial ponds along each of three parallel transects 30m, 120m, and 480m from a source wetland consisting of a stream (Riacho Passarinho) and its floodplain. Within each transect, the distance between adjacent artificial ponds was 30 m. The well-drained sandy soils ensured that no other ponds and puddles formed during the rainy season at our study site, which could confound our manipulation of isolation distances. Each fish-by-distance treatment was replicated four times for a total of 24 artificial ponds.

Our experiment ran from 18-Jan-2017 to 24-Apr-2017 and was intended to simulate the seasonal dynamics of temporary or semi-permanent ponds, which experience drought every dry season. Such ponds are filled with water during heavy rains and dry out slowly for the next few months (e.g. da Silva et al. 2012, Schiesari and Corrêa 2016) being largely recolonized by macroinvertebrates every rainy season and our experiments should be interpreted as applying to understand the dynamics of a single such colonization cycle. Temporary and semi-permanent ponds only rarely contain fish and any differences we find with respect to fish effects are of limited direct applicability even though they may be the role of important mechanisms of interaction and dynamics with more general application.

Between 18 and 25-Jan-2017 mesocosms were filled with well water. Fish were added on 29-Jan-2017. We conducted three sampling surveys of freshwater macroinvertebrates after ∼3 weeks (18 to 23-Feb-2017), ∼8 weeks (23 to 27-Mar-2017), and ∼12 weeks (20 to 24-Apr-2017) of the experiment. We have previously used data from this experiment to address other questions about community structure and more detailed information on the experimental design and data collection is available in the Methods and Supplementary Material of Pelinson et al. (2020).

### Data analysis

In the rest of this paper, we focused on the second and third surveys. We ignored the first survey (∼3 weeks) because we found that stochasticity was very high as might be predicted (but see appendix S3 for details). We tested for the effect of isolation, presence of fish, survey and their interactions on the total abundance of individuals (*i*.*e*., community size) and observed richness through type II Wald chi-square tests for generalized linear mixed models. Total abundances were modeled using a negative binomial distribution due to overdispersion, whereas richness was modeled using a normal distribution. The identity of ponds was considered a random effect term in the models. We also performed *posthoc* pairwise comparisons to assess differences among specific treatments adjusting p-values for multiple comparisons using the Šidák method. We are lacking data from four ponds in the third survey due to logistical issues; therefore, in the third survey, treatments with fish in 30 m, 120 m, and 480 m, and without fish in 480 m, had only three replicates (see Pelinson *et al*. 2020 for details). We compared gamma diversity in each treatment by computing the effective number of species (*i*.*e*., Hill numbers q = 0, Chao *et al*. 2014; Hsieh *et al*. 2016), that is, we computed the number of species in each treatment for a similar number of sampled ponds through sample-based rarefaction and extrapolation (Colwell et al. 2012). Note here that what we call gamma-diversity is the realized species pool that was able to colonize a given treatment.

To test how within-treatment beta-diversity is affected by isolation and presence of fish, we used the distance of each replicate pond (*i*.*e*., communities) to their group spatial centroid in multivariate space as response variables (Anderson et al. 2006). This is achieved by placing a distance matrix of any measure of dissimilarity between pairs of observations into a multivariate Euclidean space through principal coordinate analysis (PCoA; Anderson *et al*. 2006). In this case, greater mean distances to the treatment group spatial centroid reflect larger beta-diversity. We used the Bray-Curtis dissimilarity index on raw abundance data to compute the dissimilarity matrices. To tease apart the community dissimilarity caused by variations in community size, gamma, and alpha diversity from deterministic processes leading to homogenization or divergence of structures, we used a null model approach to calculate the expected community similarity in the absence of such processes in 1,000 simulated communities (Chase et al. 2011). We used a null model that shuffles individuals across communities but preserves the number of absences in the matrix (*i*.*e*., matrix fill), total species abundances in each community (*i*.*e*., row sums), and species abundances distributions in the metacommunity (*i*.*e*., column sums) for each treatment separately to keep gamma and average alpha diversity constant in each treatment. By not shuffling individuals among treatments, we were able to assess the expected stochasticity in a landscape that supposedly only contains ponds of the specific condition described by its assigned treatment. Then, for each community we calculated how much distance to its group centroid deviates from the distance that would be expected by the null model (*i*.*e*., beta-deviation; Chase *et al*. 2011; Kraft *et al*. 2011) using:

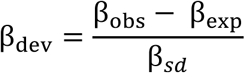

where β_obs_ is the observed distance to the centroid, β_exp_ is the mean distance to centroid expected by the null model, and β_*sd*_ is the standard deviation of the expected distances. This method enables statistical tests of treatment effects on both mean observed beta-diversity and mean beta-deviation (see Vannette & Fukami 2017). The mean observed beta-diversity of a given treatment is a measure of community variability, the higher the observed mean beta-diversity, the more variable communities are. The mean beta-deviation, however, is a measure of the non-random effect of a treatment on community variability (Xing and He 2020).

This null model approach has been questioned by some authors for not accurately removing the effect of differences in alpha and gamma diversity from beta-diversity (Ulrich et al. 2017, Xing and He 2020). Note, however, that this is not our goal here. We computed beta-deviation as a relative rather than an absolute estimate of how much community variability within treatments can be attributed to stochastic and non-stochastic processes. Another shortcoming of the beta-deviation approach is that, as it is not fully independent from gamma, it tends to scale with sampling effort (Bennett and Gilbert 2016, Xing and He 2020). Even though we had four out of 12 treatments with three replicates instead of four (see details in Pelinson *et al*. 2020), we did not observe significant differences in gamma diversity among treatments, therefore it is unlikely to qualitatively affect our results.

We tested for differences of observed distances to centroid and beta-deviations among treatments also using linear mixed models with treatments as fixed terms and pond identity as a random effect term. The effect of treatments was tested through type II Wald chi-square tests. We also did *posthoc* pairwise comparisons adjusting p-values using the Šidák method. All analyses were performed in R version 4.0.2 (R Core Team 2020). Sample-based rarefaction and extrapolations were done using the package ‘iNEXT’ version 2.0.19 (Hsieh et al. 2016). The null communities were generated using the “quasiswap” algorithm from the package ‘vegan’ version 2.5-6 (Oksanen et al. 2019). All code used to analyze data and reproduce figures is available in the GitHub repository: RodolfoPelinson/PredatorIsolationStochasticity.

## RESULTS

Gamma diversity (*i*.*e*., treatment richness) was generally not different among treatments, except for fishless ponds in the intermediate isolation treatment in the last survey, which had the highest number of species in our experiment (Fig. 2a). Community size, however, significantly grew from the second to the third survey, and this effect was particularly strong in more isolated ponds, and for fishless ponds (Table 1; Fig. 2b).

**Table 1.** Deviance table of type II Wald Chi-square tests for the generalized linear mixed models. Only significant differences in *Post-hoc* comparisons are shown.

|  | Df | Chi-square | p | *Post-hoc* comparisons |
| --- | --- | --- | --- | --- |
| *Abundance* |  |  |  |  |
| Fish | 1 | 0.013 | 0.909 |  |
| Isolation | 2 | 0.323 | 0.851 |  |
| **Sampling Survey** | **1** | **37.225** | **<0.001** |  |
| Fish : Isolation | 2 | 1.609 | 0.447 |  |
| **Fish : Sampling Survey** | **1** | **4.340** | **0.037** | Fishless: 2nd << 3rd; Fish: 2nd < 3rd |
| **Isolation : Sampling Survey** | **2** | **12.689** | **0.002** | 120m: 2nd < 3rd; 480m: 2nd << 3rd |
| Fish : Isolation : Sampling Survey | 2 | 0.597 | 0.742 |  |
| *Observed Beta-diversity* |  |  |  |  |
| Fish | 1 | 1.442 | 0.230 |  |
| Isolation | 2 | 2.071 | 0.355 |  |
| Sampling Survey | 1 | 2.136 | 0.144 |  |
| **Fish : Isolation** | **2** | **17.217** | **<0.001** | Fish: 30m > 480m |
| Fish : Sampling Survey | 1 | 0.086 | 0.769 |  |
| **Isolation : Sampling Survey** | **2** | **7.317** | **0.026** |  |
| Fish : Isolation : Sampling Survey | 2 | 5.463 | 0.065 |  |
| *Beta-deviation* |  |  |  |  |
| **Fish** | **1** | **4.129** | **0.042** |  |
| Isolation | 2 | 4.739 | 0.094 |  |
| Sampling Survey | 1 | 1.048 | 0.306 |  |
| **Fish : Isolation** | **2** | **9.345** | **0.009** | Fishless: 30m < 120m, 30m < 480m |
| Fish : Sampling Survey | 1 | 0.096 | 0.757 |  |
| Isolation : Sampling Survey | 2 | 2.445 | 0.295 |  |
| Fish : Isolation : Sampling Survey | 2 | 2.595 | 0.273 |  |

**Figure 2.**
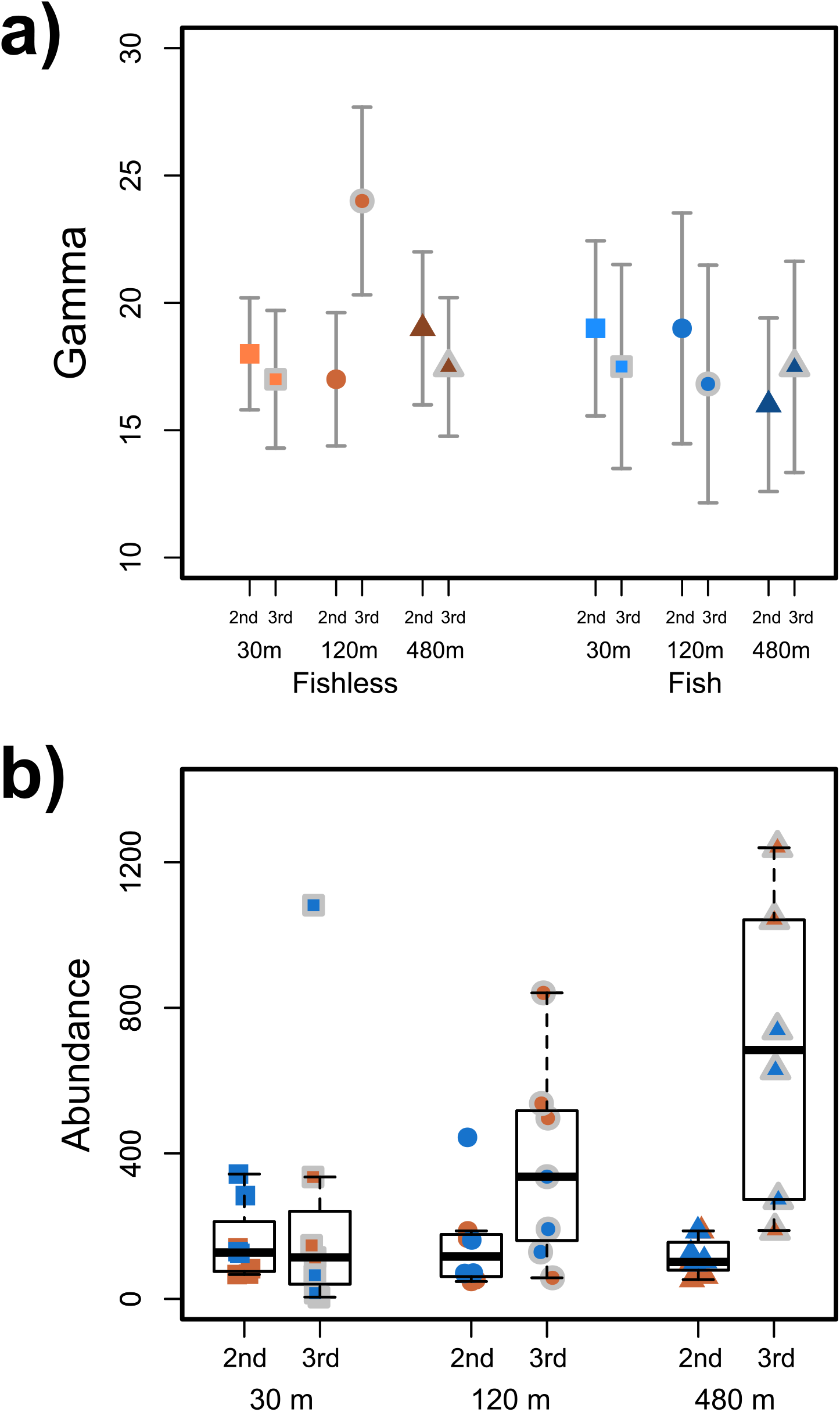
(a) Gamma diversity (regional richness) per treatment in the second and third surveys considering four sample units per treatment. Grey bars are estimated 95% confidence intervals. Estimates which confidence intervals do not overlap are significantly different from each other. (b) Box plot of abundance (*i*.*e*., community size) by sampling survey in each isolation treatment. Squares, circles, and triangles are respectively 30 m, 120 m, and 480 m isolation treatments. Orange represents fishless ponds, whereas blue represents ponds with fish. Symbols without a grey border are from the second survey, whereas symbols with a gray border are from the third survey.

Observed beta-diversity slightly increased from the second to the third survey, but only in low isolation. Both changes in observed beta-diversity and beta-deviation were significantly explained by an interaction between spatial isolation and the presence of fish (Table 1). Observed beta-diversity did not significantly change across the isolation gradient in fishless ponds (Fig. 3a). However, the values of beta-deviation for fishless ponds significantly increased from 30m to 120 and 480m of isolation (Table 1; Fig. 3b). Additionally, values of beta-deviation for these ponds went from values close to 0 in 30m isolation to positive values in 120m and 480m of isolation, so that beta-diversity was higher than expected by null models in more isolated ponds. When we considered only ponds with fish, we observed a significant decrease in observed beta-diversity from 30m to 480m of isolation (Table 1; Fig. 3a). More importantly, values of beta-deviation for ponds with fish did not differ significantly among isolation treatments with values always close to the 0 line (*i*.*e*. observed beta-diversity always similar to expected by the null models; Fig. 3b).

**Figure 3.**
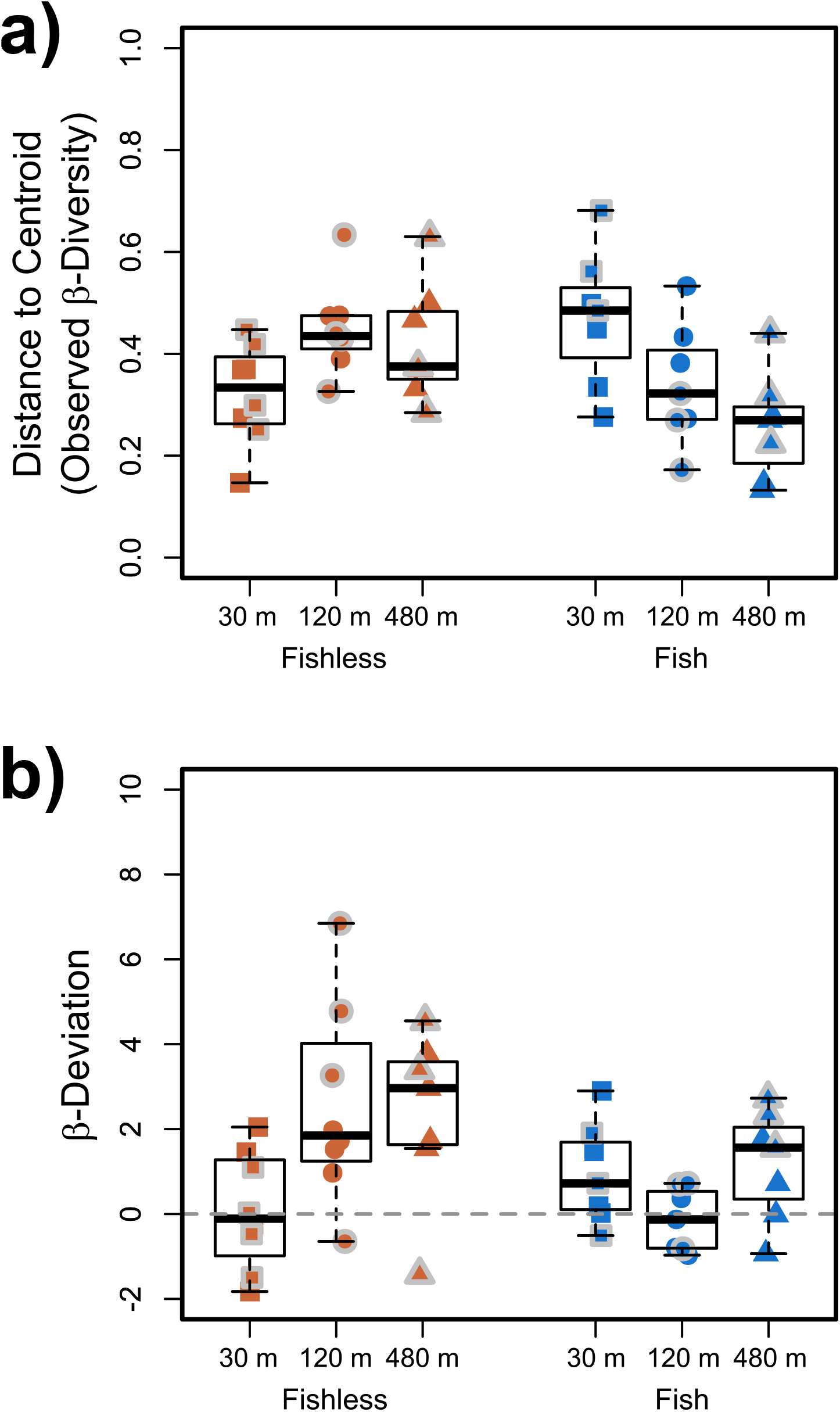
Box plots of values of distance to the centroid of observed dissimilarity values based on Bray-Curtis distance (a) and distance to centroid based on beta-deviation measures (b). Squares, circles, and triangles are 30 m, 120 m, and 480 m isolation treatments, respectively. Orange squares, circles, and triangles are fishless ponds, whereas blues are ponds with fish. Squares, circles, and triangles without a grey border are from the second survey, whereas the ones with a gray border are from the third survey.

## DISCUSSION

Few studies have tried to separate the contributions of stochastic vs environment-independent non-stochastic assembly processes on composition change (*i*.*e*. beta-diversity or community variability) (*e*.*g*., Vannette and Fukami 2017). Because our experimental and analytical design controlled for environmental heterogeneity and dispersal limitation among experimental mesocosms, community variability could be attributed to stochasticity in dispersal and/or demography or species interactions (*e*.*g*. priority effect). Because we equally accounted for stochasticity in all treatments by using null models, the importance of deterministic processes affecting community variability, such as priority effects and density-dependent population growth, were manifest as beta-deviation. Therefore, even though our null scenario may carry some deterministic processes that generated the original community and metacommunity structures, here we show relative contributions of stochastic and non-stochastic assembly processes to community variability in different scenarios of dispersal limitation, and the presence or absence of a generalist top predator.

We predicted that predatory fish would decrease community variability and decrease gamma diversity because they can exclude a defined subset of vulnerable taxa(H1; Chase *et al*. 2009). Contrary to this expectation, however, we found that fish had no significant overall effects on mean gamma or beta-diversity in our experiment. We also expected that the negative effects of fish on insect gamma diversity would be stronger for predatory insects than for insects at lower trophic levels since predatory insects tend to be more vulnerable to visually oriented predators (Diehl 1992, Wellborn et al. 1996, Batzer et al. 2000, Boelter et al. 2018). However, even though we found that fish did have a stronger negative effect on the abundance of predatory insects when compared to herbivores and detritivores (see appendix S6), they did not regionally exclude species from ponds. Tilapias are known as generalist omnivorous fish, and although they might preferentially prey upon predatory insects because they are easier to see, it is likely that their prey preference would shift according to prey availability. Therefore, they could thus effectively decrease predatory insect abundances, but not regional richness.

We also predicted that community variability would decrease with spatial isolation because unequal dispersal rates would decrease gamma diversity (H2). We found that community variability did decrease with isolation in ponds with fish, but this decrease was not present after accounting for stochastic effects of demographic drift and chance dispersal with null models. We also found that dispersal limitation was not sufficient to completely exclude poor dispersers from isolated ponds. Rather, the decrease in beta-diversity was likely due to an increase in community size with isolation. Other work has shown that because herbivores and detritivores are less dispersal limited than predatory insects, their populations tend to increase in more isolated habitats due to a phenomenon analogous to a trophic cascade (Shulman and Chase 2007, Chase and Shulman 2009, Pelinson et al. 2020). Therefore, here we observe that the increase in abundance of herbivores and detritivores caused a general increase in community size, likely decreasing the importance of ecological drift on community structure, thus decreasing community variability along the isolation gradient (Fig. 4a).

**Figure 4.**
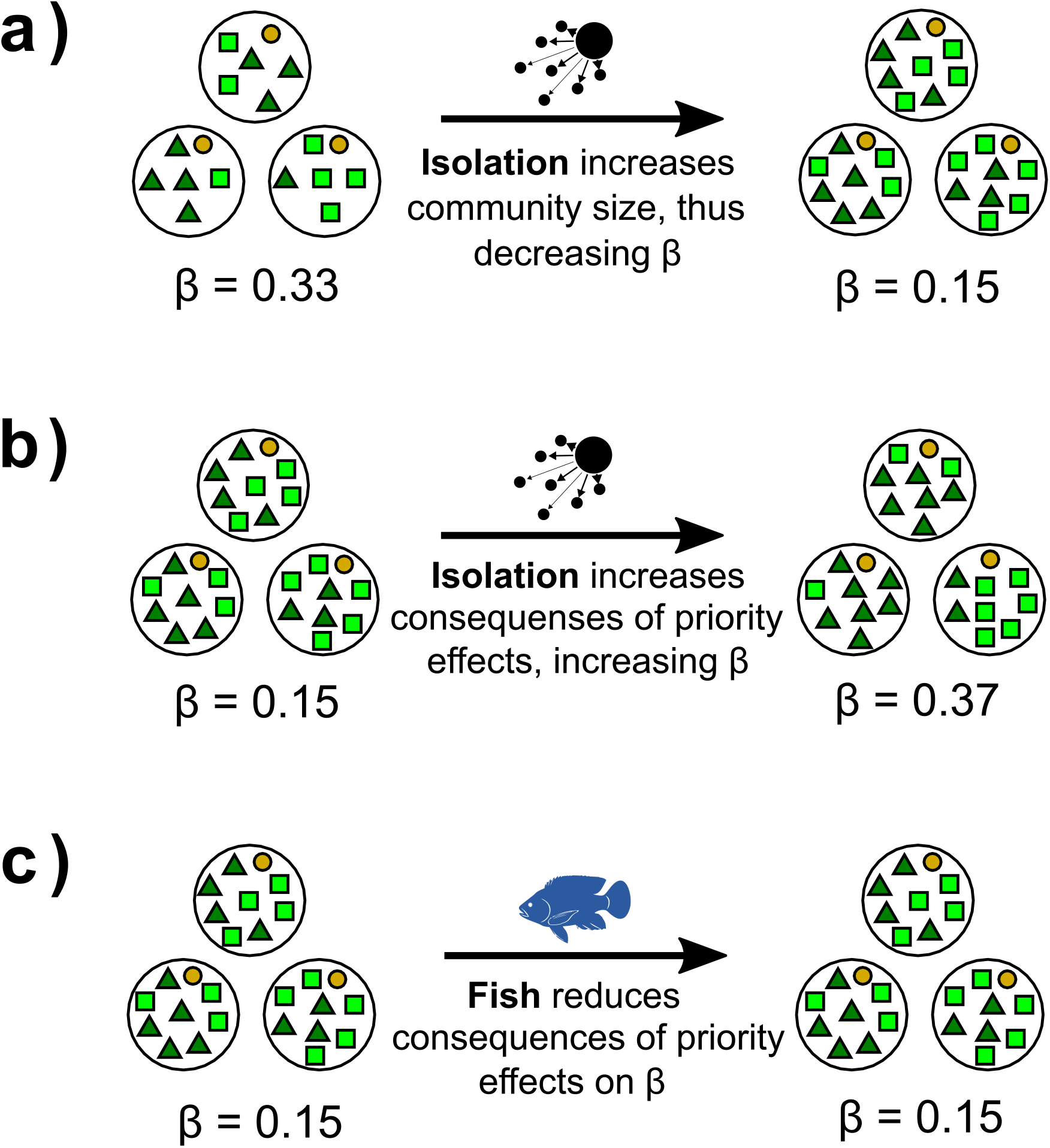
We found that: (a) spatial isolation can decrease community variability by increasing community size due to effects similar to trophic cascades, thus reducing the importance of ecological drift on community structure; (b) Spatial isolation may also increase community variability by increasing the importance of deterministic processes not related to environmental heterogeneity, possibly priority effects, on community assembly making communities more variable them would be expected in the absence of such processes; (c) however, when fish is present, it can reduce the importance of these priority effects.

In contrast, we found no effect of isolation on observed community variability for fishless ponds, even though the increase in community size also happened in those ponds. Most of the evidence and theoretical models of single trophic level metacommunities points out to a positive effect of dispersal limitation on community variability (Mouquet and Loreau 2003, Astorga et al. 2012, Grainger and Gilbert 2016). This effect is, in general, due to either the decrease in the local richness that low dispersal confers (Mouquet and Loreau 2003, Chase and Myers 2011, Leibold and Chase 2018) (see Appendix S1), which we did not observe (see appendix S4), or to an increase in the importance of priority effects when dispersal rates are similar among species (see Appendix S1) (Vellend et al. 2014). Because we know that some freshwater insects are more abundant than others in more isolated ponds (Shulman and Chase 2007, Chase and Shulman 2009), we assumed that they vary in dispersal rates. Thus, we expected priority effects to decrease beta-diversity because the order of arrival of species in isolated ponds would be determined by dispersal frequency (H3; Vellend *et al*. 2014). However, most of those differences are correlated with trophic level (Shulman and Chase 2007, Chase and Shulman 2009), and, because of the numerical dominance of herbivores and detritivores in our experiment (85% of the individuals and 66% of the taxa), it is plausible to assume that dispersal rates of most species were similar enough to make the order of the first colonizers sufficiently random, even in highly isolated habitats. In this scenario, priority effects could have led communities to different structures (Vellend et al. 2014, Fukami 2015). In support of this hypothesis, we observed that the values of beta-deviation in fishless ponds increased with spatial isolation, meaning that communities became more different from each other than what would be expected by random chance (Fig. 4b). Therefore, we argue that the lack of effect of isolation on community variability in fishless ponds is due to the net consequences of the increase in community size (Fig. 4a), which makes communities less variable, and an increase in the importance of priority effects, which makes communities more variable (Fig. 4b).

Priority effects are likely to play an important role in community variability when organisms have rapid life cycles, thus being able to establish large populations and resist invasion by species arriving later (Louette and De Meester 2007, Leopold et al. 2017, Vannette and Fukami 2017). However, most freshwater insects inhabiting temporary ponds spend only part of their life cycle in water, making a demographic preemption, as mentioned above, unlikely. Indeed, competition has been shown to affect freshwater insect and tadpole biomass and time to metamorphose, but not abundance patterns (Blaustein and Margalit 1994, 1996). Alternatively, priority effects in this system could arise due to other post or pre-colonization species interactions. Predation (including intraguild predation), for instance, is common among freshwater insects (*e*.*g*., Wissinger and McGrady 1993, Wissinger et al. 1996, Fincke 1999). In this scenario, individuals arriving first can become large enough to prey upon any other insects arriving later. Pre-colonization habitat selection could also yield similar consequences in community structure (Fukami 2015). Numerous works have shown that colonization by many aquatic animals is behaviorally dependent on the identity and density of species that have already colonized a habitat (*e*.*g*., Resetarits 2005, Sadeh et al. 2009, Kraus and Vonesh 2010, Pintar and Resetarits 2017, Trekels and Vanschoenwinkel 2019). For instance, several different species of mosquitoes have been shown to detect and avoid oviposition in ponds with both competitors (Blaustein and Kotler 1993) and predators (backswimmers: Eitam et al. 2002, Kiflawi et al. 2003, Blaustein et al. 2004, Negrão 2019; dragonflies: Stav et al. 1999; Negrão 2019; a combination of predators: Negrão 2019). Therefore, the infrequent arrival in more isolated habitats could have strong and fast consequences on the choice of oviposition of insects arriving later, leading more isolated habitats to be more variable than would be expected in the absence of such constraints.

We did not observe that priority effects led to higher community variability in ponds with fish, as expected (H4). We expected that fish would reduce priority effects on community structure by preying upon the most abundant and vulnerable species, thus minimizing the advantage of arriving first. We know that predatory fish had a strong negative effect on predatory insects in this experiment (Pelinson et al. 2020). We also know that fish had a negative effect on herbivores and detritivores, counteracting the indirect positive effect that the absence of predatory insects would have on them (Pelinson et al. 2020). Thus, fish probably prevented both predatory and non-predatory insects from becoming more abundant, irrespective of the order of arrival. However, we do not know to what degree these effects were a result of direct predation by fish, habitat selection, or both. Fish can strongly affect the chance of oviposition of aquatic insects, regardless of their trophic level (Vonesh et al. 2009). Therefore, because fish were uniformly present in ponds treated with fish, the habitat selection responses of aquatic insects might have pushed communities toward more similar community structures. Even though we cannot precise by what mechanism, we nevertheless show that the presence of fish can override deterministic effects caused by factors other than environmental heterogeneity on community structure (Fig. 4c).

Here we show that dispersal limitation can have both negative and positive effects on community variability in freshwater insect communities. Negative effects are likely because the unequal dispersal rates among predatory insects and their prey cause community size to increase with isolation due to trophic cascades (Shulman and Chase 2007, Chase and Shulman 2009, Hein and Gillooly 2011, Pelinson et al. 2020). Such an increase in community size can negatively affect community variability by reducing the consequences of demographic stochasticity on community structure. Positive effects, however, are likely because of an increase in the importance of priority effects in generating different community structures, even though we could not identify if it is due to pre (*i*.*e*., habitat selection) or post-colonization (*i*.*e*., predation and competition) mechanisms. Due to the short duration of our experiment, our results for fishless habitats are mostly valid for temporary ponds. Longer experiments (*i*.*e*. more than one year) would be necessary to evaluate the validity of these results for permanent fishless ponds.

We also show that, when fish are present, priority effects are diminished, causing community variability to decrease along the isolation gradient due to the increase in community size. We believe this result can be extrapolated to act in more permanent ponds since its main driver is the increase in the abundance of herbivores and detritivores, which only became stronger with time, and have been observed in other experiments and natural habitats (Shulman and Chase 2007, Chase and Shulman 2009). Finally, our findings show that considering multiple trophic levels can be important in predicting patterns of community variability since inter-trophic level effects can substantially change how different processes affect community size and priority effects, and consequently, community variability.

## ACKNOWLEDGMENTS

We thank the EESB staff for assistance in pond construction and Luis Vicente P. Cavalaro, Bianca S. Valente, Fernanda Simioni, Débora Negrão, Jessika Akane and Suzana Marte for assistance in the community sampling surveys. We thank Tadeu Siqueira and Paulo Inácio Prado for conceptual and statistical advice, Victor Saito and Erika Shimabukuro for assistance in the identification of aquatic insects, Renata Pardini and Daniel Lahr for providing lab and office space, Paulo Roberto Guimarães Junior, Paulo De Marco Júnior and Jonathan Chase for insightful comments on early versions of the manuscript; The members of the Schiesari lab and Leibold lab for criticism; and Giselda Durigan for introducing us to the EESB. This study was funded by Fundação de Amparo à Pesquisa do Estado de São Paulo (FAPESP, grant #2014/16320-7, Carlos Navas PI, Luis Schiesari Associate Researcher; FAPESP, grant #2015/18790-3; Luiz Antonio Martinelli PI, Luis Schiesari Co-PI) and Conselho Nacional de Desenvolvimento Científico e Tecnológico (CNPq, grant #458796/2014-0). RMP was supported by Ph.D. fellowships from FAPESP (grants #2017/04122-4 and #2018/07714-2) and Coordenação de Aperfeiçoamento de Pessoal de Nível Superior (CAPES). Experiments were conducted in EESB under the authorization from Instituto Florestal (COTEC 553/2017) and collection permits from Instituto Chico Mendes de Conservação da Biodiversidade (ICMBIo 17559-6), following protocols approved by the Research Ethics Committee of the School of Arts, Sciences and Humanities of the University of São Paulo (CEUA 003/2016).

## REFERENCES

Anderson, M. J., T. O. Crist, J. M. Chase, M. Vellend, B. D. Inouye, A. L. Freestone, N. J. Sanders, H. V. Cornell, L. S. Comita, K. F. Davies, S. P. Harrison, N. J. B. Kraft, J. C. Stegen and N. G. Swenson. 2011. Navigating the multiple meanings of β diversity: a roadmap for the practicing ecologist. Ecology Letters 14:19–28.

Anderson, M. J., K. E. Ellingsen, and B. H. McArdle. 2006. Multivariate dispersion as a measure of beta diversity. Ecology Letters 9:683–693.

Astorga, A., J. Oksanen, M. Luoto, J. Soininen, R. Virtanen, and T. Muotka. 2012. Distance decay of similarity in freshwater communities: do macro- and microorganisms follow the same rules? Global Ecology and Biogeography 21:365–375.

Barwell, L. J., N. J. B. Isaac, and W. E. Kunin. 2015. Measuring β-diversity with species abundance data. Journal of Animal Ecology 84:1112–1122.

Batzer, D. P., C. R. Pusateri, and R. Vetter. 2000. Impacts of fish predation on marsh invertebrates: Direct and indirect effects. Wetlands 20:307–312.

Bennett, J. R., and B. Gilbert. 2016. Contrasting beta diversity among regions: how do classical and multivariate approaches compare? Global Ecology and Biogeography 25:368–377.

Blaustein, L., M. Kiflawi, A. Eitam, M. Mangel, and J. E. Cohen. 2004. Oviposition habitat selection in response to risk of predation in temporary pools: mode of detection and consistency across experimental venue. Oecologia 138:300–305.

Blaustein, L., and B. P. Kotler. 1993. Oviposition habitat selection by the mosquito, Culiseta longiareolata: effects of conspecifics, food and green toad tadpoles. Ecological Entomology 18:104–108.

Blaustein, L., and J. Margalit. 1994. Mosquito larvae (Culiseta longiareolata) prey upon and compete with toad tadpoles (Bufo viridis). Journal of Animal Ecology 63:841–850.

Blaustein, L., and J. Margalit. 1996. Priority effects in temporary pools: nature and outcome of mosquito larva-toad tadpole interactions depend on order of entrance. Journal of Animal Ecology 65:77–84.

Boelter, T., C. Stenert, M. M. Pires, E. S. F. Medeiros, and L. Maltchik. 2018. Influence of plant habitat types and the presence of fish predators on macroinvertebrate assemblages in southern Brazilian highland wetlands. Fundamental and Applied Limnology 192:65–77.

Chao, A., N. J. Gotelli, T. C. Hsieh, E. L. Sander, K. H. Ma, R. K. Colwell, and A. M. Ellison. 2014. Rarefaction and extrapolation with Hill numbers: a framework for sampling and estimation in species diversity studies. Ecological Monographs 84:45–67.

Chase, J. M. 2003. Community assembly: when should history matter? Oecologia 136:489–498.

Chase, J. M. 2007. Drought mediates the importance of stochastic community assembly. Proceedings of the National Academy of Sciences 104:17430–17434.

Chase, J. M. 2010. Stochastic community assembly causes higher biodiversity in more productive environments. Science 328:1388–1391.

Chase, J. M., E. G. Biro, W. A. Ryberg, and K. G. Smith. 2009. Predators temper the relative importance of stochastic processes in the assembly of prey metacommunities. Ecology Letters 12:1210–1218.

Chase, J. M., N. J. B. Kraft, K. G. Smith, M. Vellend, and B. D. Inouye. 2011. Using null models to disentangle variation in community dissimilarity from variation in α-diversity. Ecosphere 2:art24.

Chase, J. M., and J. A. Myers. 2011. Disentangling the importance of ecological niches from stochastic processes across scales. Philosophical Transactions of the Royal Society B: Biological Sciences 366:2351–2363.

Chase, J. M., and R. S. Shulman. 2009. Wetland isolation facilitates larval mosquito density through the reduction of predators. Ecological Entomology 34:741–747.

Colwell, R. K., A. Chao, N. J. Gotelli, S.-Y. Lin, C. X. Mao, R. L. Chazdon, and J. T. Longino. 2012. Models and estimators linking individual-based and sample-based rarefaction, extrapolation and comparison of assemblages. Journal of Plant Ecology 5:3–21.

Connor, E. F., and D. Simberloff. 1979. The assembly of species communities: chance or competition? Ecology 60:1132–1140.

Diamond, J. M. 1975. “Assembly of species communities. Pages 342–444 in M. L. Cody and J. M. Diamond, editors. Ecology and evolution of communities. The Belknap Press of Harvard University Press, Cambridge, Massachusetts, and London, England.

Diehl, S. 1992. Fish predation and benthic community structure: the role of omnivory and habitat complexity. Ecology 73:1646–1661.

Eitam, A., L. Blaustein, and M. Mangel. 2002. Effects of Anisops sardea (Hemiptera: Notonectidae) on oviposition habitat selection by mosquitoes and other dipterans and on community structure in artificial pools. Hydrobiologia 485:183–189.

Fincke, O. M. 1999. Organization of predator assemblages in Neotropical tree holes: effects of abiotic factors and priority. Ecological Entomology 24:13–23.

Fukami, T. 2015. Historical contingency in community assembly: integrating niches, species pools, and priority effects. Annual Review of Ecology, Evolution, and Systematics 46:1– 23.

Goyke, A. P., and A. E. Hershey. 1992. Effects of fish predation on larval chironomid (Diptera: Chironomidae) communities in an arctic ecosystem. Hydrobiologia 240:203–211.

Grainger, T. N., and B. Gilbert. 2016. Dispersal and diversity in experimental metacommunities: linking theory and practice. Oikos 125:1213–1223.

Hein, A. M., and J. F. Gillooly. 2011. Predators, prey, and transient states in the assembly of spatially structured communities. Ecology 92:549–555.

Hendrickx, F., J.-P. Maelfait, K. Desender, S. Aviron, D. Bailey, T. Diekotter, L. Lens, J. Liira, O. Schweiger, M. Speelmans, V. Vandomme, and R. Bugter. 2009. Pervasive effects of dispersal limitation on within- and among-community species richness in agricultural landscapes. Global Ecology and Biogeography 18:607–616.

Hijmans, R. J., S. E. Cameron, J. L. Parra, P. G. Jones, and A. Jarvis. 2005. Very high resolution interpolated climate surfaces for global land areas. International Journal of Climatology 25:1965–1978.

Hsieh, T. C., K. H. Ma, and A. Chao. 2016. iNEXT: an R package for rarefaction and extrapolation of species diversity (Hill numbers). Methods in Ecology and Evolution 7:1451–1456.

Hubbell, S. P. 2001. The unified neutral theory of biodiversity and biogeography. Princeton University Press.

Kiflawi, M., L. Blaustein, and M. Mangel. 2003. Predation-dependent oviposition habitat selection by the mosquito Culiseta longiareolata: a test of competing hypotheses. Ecology Letters 6:35–40.

Koleff, P., K. J. Gaston, and J. J. Lennon. 2003. Measuring beta diversity for presence–absence data. Journal of Animal Ecology 72:367–382.

Kottek, M., J. Grieser, C. Beck, B. Rudolf, and F. Rubel. 2006. World Map of the Köppen-Geiger climate classification updated. Meteorologische Zeitschrift:259–263.

Kraft, N. J. B., L. S. Comita, J. M. Chase, N. J. Sanders, N. G. Swenson, T. O. Crist, J. C. Stegen, M. Vellend, B. Boyle, M. J. Anderson, H. V. Cornell, K. F. Davies, A. L. Freestone, B. D. Inouye, S. P. Harrison, and J. A. Myers. 2011. Disentangling the drivers of β diversity along latitudinal and elevational gradients. Science 333:1755–1758.

Kraus, J. M., and J. R. Vonesh. 2010. Feedbacks between community assembly and habitat selection shape variation in local colonization. Journal of Animal Ecology 79:795–802.

Law, R., and R. D. Morton. 1993. Alternative permanent states of ecological communities. Ecology 74:1347–1361.

Leibold, M. A. 1996. A Graphical Model of Keystone Predators in Food Webs: Trophic regulation of abundance, incidence, and diversity patterns in communities. The American Naturalist 147:784–812.

Leibold, M. A. 1999. Biodiversity and nutrient enrichment in pond plankton communities. Evolutionary Ecology Research 1:73–95.

Leibold, M. A., and J. M. Chase. 2018. Metacommunity Ecology. Princeton University Press, Princeton, New Jersey.

Leopold, D. R., J. P. Wilkie, I. A. Dickie, R. B. Allen, P. K. Buchanan, and T. Fukami. 2017. Priority effects are interactively regulated by top-down and bottom-up forces: evidence from wood decomposer communities. Ecology Letters 20:1054–1063.

Louette, G., and L. De Meester. 2007. Predation and priority effects in experimental zooplankton communities. Oikos 116:419–426.

Melo, A. C. G., and G. Durigan. 2011. Estação Ecológica de Santa Bárbara Plano de Manejo. Secretaria do Meio Ambiente. Secretaria do Meio Ambiente.

Mouquet, N., and M. Loreau. 2003. Community patterns in source-sink metacommunities. The American Naturalist 162:544–557.

Myers, J. A., J. M. Chase, R. M. Crandall, and I. Jiménez. 2015. Disturbance alters beta-diversity but not the relative importance of community assembly mechanisms. Journal of Ecology 103:1291–1299.

Negrão, D. S. G. 2019. Contágio espacial resultante do risco de predação na seleção de sí•tios de oviposição por mosquito. Thesis, Universidade de São Paulo, São Paulo.

Oksanen, J., F. G. Blanchet, M. Friendly, R. Kindt, P. Legendre, D. McGlinn, P. R. Minchin, R. B. O’Hara, G. L. Simpson, P. Solymos, M. H. H. Stevens, E. Szoecs, and H. Wagner. 2019. vegan: Community Ecology Package.

Orrock, J. L., and R. J. Fletcher Jr. 2005. Changes in community size affect the outcome of competition. The American Naturalist 166:107–111.

Pelinson, R. M., M. A. Leibold, and L. Schiesari. 2020. Top predator introduction changes the effects of spatial isolation on freshwater community structure. bioRxiv:857318.

Pintar, M. R., and W. J. Resetarits. 2017. Context-dependent colonization dynamics: Regional reward contagion drives local compression in aquatic beetles. Journal of Animal Ecology 86:1124–1135.

R Core Team. 2020. R: A language and environment for statistical computing. Page R Foundation for Statistical Computing. Vienna, Austria.

Resetarits, W. J. 2005. Habitat selection behaviour links local and regional scales in aquatic systems. Ecology Letters 8:480–486.

Sadeh, A., M. Mangel, and L. Blaustein. 2009. Context-dependent reproductive habitat selection: the interactive roles of structural complexity and cannibalistic conspecifics. Ecology Letters 12:1158–1164.

Schiesari, L., and D. T. Corrêa. 2016. Consequences of agroindustrial sugarcane production to freshwater biodiversity. GCB Bioenergy 8:644–657.

Shulman, R. S., and J. M. Chase. 2007. Increasing isolation reduces predator:prey species richness ratios in aquatic food webs. Oikos 116:1581–1587.

Shurin, J. B., P. Amarasekare, J. M. Chase, R. D. Holt, M. F. Hoopes, and M. A. Leibold. 2004. Alternative stable states and regional community structure. Journal of Theoretical Biology 227:359–368.

da Silva, F. R., C. P. Candeira, and D. de C. Rossa-Feres. 2012. Dependence of anuran diversity on environmental descriptors in farmland ponds. Biodiversity and Conservation 21:1411– 1424.

Siqueira, T., V. S. Saito, L. M. Bini, A. S. Melo, D. K. Petsch, V. L. Landeiro, K. T. Tolonen, J. Jyrkänkallio-Mikkola, J. Soininen, and J. Heino. 2020. Community size can affect the signals of ecological drift and niche selection on biodiversity. Ecology 101:e03014.

Stav, G., L. Blaustein, and J. Margalith. 1999. Experimental evidence for predation risk sensitive oviposition by a mosquito, Culiseta longiareolata. Ecological Entomology 24:202–207.

Trekels, H., and B. Vanschoenwinkel. 2019. Both local presence and regional distribution of predator cues modulate prey colonisation in pond landscapes. Ecology Letters 22:89–97.

Ulrich, W., A. Baselga, B. Kusumoto, T. Shiono, H. Tuomisto, and Y. Kubota. 2017. The tangled link between β- and γ-diversity: a Narcissus effect weakens statistical inferences in null model analyses of diversity patterns. Global Ecology and Biogeography 26:1–5.

Vannette, R. L., and T. Fukami. 2017. Dispersal enhances beta diversity in nectar microbes. Ecology Letters 20:901–910.

Vellend, M. 2016. The Theory of Ecological Communities. Princeton University Press, Princeton.

Vellend, M., D. S. Srivastava, K. M. Anderson, C. D. Brown, J. E. Jankowski, E. J. Kleynhans, N. J. B. Kraft, A. D. Letaw, A. A. M. Macdonald, J. E. Maclean, I. H. Myers-Smith, A. R. Norris, and X. Xue. 2014. Assessing the relative importance of neutral stochasticity in ecological communities. Oikos 123:1420–1430.

Vonesh, J. R., J. M. Kraus, J. S. Rosenberg, and J. M. Chase. 2009. Predator effects on aquatic community assembly: disentangling the roles of habitat selection and post-colonization processes. Oikos 118:1219–1229.

Wellborn, G. A., D. K. Skelly, and E. E. Werner. 1996. Mechanisms creating community structure across a freshwater habitat gradient. Annual Review of Ecology and Systematics 27:337–363.

Whittaker, R. H. 1960. Vegetation of the Siskiyou Mountains, Oregon and California. Ecological Monographs 30:279–338.

Whittaker, R. H. 1972. Evolution and Measurement of Species Diversity. Taxon 21:213–251.

Wissinger, S. A., G. B. Sparks, G. L. Rouse, W. S. Brown, and H. Steltzer. 1996. Intraguild predation and Cannibalism among larvae of detritivorous caddisflies in subalpine wetlands. Ecology 77:2421–2430.

Wissinger, S., and J. McGrady. 1993. Intraguild predation and competition between larval dragonflies: direct and indirect effects on shared prey. Ecology 74:207–218.

Xing, D., and F. He. 2020. Analytical models for β-diversity and the power-law scaling of β-deviation. bioRxiv:2020.04.19.049163.

