## Appendix S1 for "Spatial community variability: Interactive effects of predators and isolation on stochastic community assembly"

### Appendix S1 – Alternative potential effects of the presence of top predators and dispersal limitation on community variability.

**Table 1.** Potential, non-mutually exclusive consequences of the presence of generalist and specialist predators on community variability via their effects on the metacommunity properties community size, alpha-diversity, and gamma-diversity.

| Type of Predator | Metacommunity aspect | Effect on metacommunity aspect | Consequence to community variability | Explanation |
|---|---|---|---|---|
| Specialist | Community Size | Negative | Positive | Specialist predators may cause a reduction in community size by preying upon the most vulnerable prey. |
| | Alpha Diversity | Negative | Positive | Constant predation pressure on a subset of vulnerable prey may reduce alpha diversity on local communities (Chase et al. 2009). |
| | | Positive | Negative | If there is a trade-off between competitive ability and vulnerability to predation, predators may act as keystone species allowing weak competitors to persist in a community (Paine 1966, Leibold 1996), thus increasing alpha diversity. |
| | Gamma Diversity | Negative | Negative | If predators preferentially prey upon a subset of more vulnerable species, those species may become regionally extinct (Chase et al. 2009). |
| Generalist | Community Size | Negative | Positive | Constant predation pressure by a generalist predator may equally decrease prey populations, decreasing community size, therefore increasing the consequences of drift on community structure (Orrock and Fletcher Jr. 2005). |
| | Alpha Diversity | Negative | Positive | Predators may cause random local extinctions by increasing the effects of ecological drift as a consequence of a reduction in community size (Ryberg and Chase 2007). |
| | Gamma Diversity | | | None expected. |

**Table 2.** Potential, non-mutually exclusive consequences of dispersal limitation in different scenarios of variation in dispersal rates on community variability via their effects on the metacommunity properties community size, alpha-diversity, and gamma-diversity.

| Dispersal limitation | metacommunity aspect | Effect on metacommunity aspect | Consequence to community variability | Explanation |
|---|---|---|---|---|
| Weak variation in Dispersal Rates | Community Size | Negative | Positive | As spatial isolation increases, the arrival of individuals from all species become equally rarer, which can reduce community size, increasing the consequences of ecological drift. |
| | Alpha Diversity | Negative | Positive | Because colonization and recolonization events become rarer, local species richness may decrease as a consequence of ecological drift (Shmida and Wilson 1985, Mouquet and Loreau 2003). |
| | Gamma Diversity | | | None expected |
| Strong variation in Dispersal Rates | Community Size | Negative | Positive | As spatial isolation increases, the arrival of more dispersal limited species become rarer, which can reduce community size, increasing the consequences of ecological drift for more dispersal limited species. |
| | Alpha Diversity | Negative | Positive | Species with lower dispersal rates can be locally excluded from more isolated habitats. |
| | Gamma Diversity | Negative | Negative | Species with lower dispersal rates can be regionally excluded from more isolated habitats (Hendrickx et al. 2009), making communities more similar to each other. |

**Table 3.** Potential, non-mutually exclusive, consequences of the presence of generalist and specialist predators, and different scenarios of variation in dispersal rates on community variability by affecting priority effects.

| | Effect on the importance of Priority effects | Consequence to community variability | Explanation |
|---|---|---|---|
| *Type of Predator* | | | |
| Specialist | Negative | Dependent on variation in dispersal | If there is a trade-off between competitive ability and vulnerability to predation, predators might only allow weak competitors to persist, decreasing the importance of priority effects (Leibold 1996, 1999, Louette and De Meester 2007). |
| Generalist | Negative | Dependent on variation in dispersal | By decreasing community size, generalist predators can reduce competition among their prey, thus decreasing the intensity of priority effects (Orrock and Fletcher Jr. 2005). |
| | Negative | Dependent on variation in dispersal | If species who arrives first become more abundant than others, and generalist predators prey mostly upon most abundant species (competitive ability/vulnerability trade-off; Leibold 1996, 1999, Louette and De Meester 2007), they will decrease the importance of priority effects. |
| | Positive | Dependent on variation in dispersal | Generalist predators can increase the importance of priority effects if their prey are strong apparent competitors (Holt et al. 1994). |
| *Dispersal limitation* | | | |
| Weak variation in Dispersal | Dependent on the strength of species interactions | Positive | Because dispersal rates among species are similar, the identity of the first colonizers is random, therefore, if species interaction are strong, priority effects will lead communities to different structures (Fukami 2015). |
| Strong variation in Dispersal | Dependent on the strength of species interactions | Negative | The identity of the first colonizers is determined by dispersal rates, therefore, if species interactions are important, it will lead communities to similar structures (Vellend et al. 2014). |

### References

Chase, J. M., E. G. Biro, W. A. Ryberg, and K. G. Smith. 2009. Predators temper the relative importance of stochastic processes in the assembly of prey metacommunities. Ecology Letters 12:1210–1218.

Fukami, T. 2015. Historical Contingency in Community Assembly: Integrating Niches, Species Pools, and Priority Effects. Annual Review of Ecology, Evolution, and Systematics 46:1–23.

Hendrickx, F., J.-P. Maelfait, K. Desender, S. Aviron, D. Bailey, T. Diekotter, L. Lens, J. Liira, O. Schweiger, M. Speelmans, V. Vandomme, and R. Bugter. 2009. Pervasive effects of dispersal limitation on within- and among-community species richness in agricultural landscapes. Global Ecology and Biogeography 18:607–616.

Holt, R. D., J. Grover, and D. Tilman. 1994. Simple Rules for Interspecific Dominance in Systems with Exploitative and Apparent Competition. The American Naturalist 144:741–771.

Leibold, M. A. 1996. A Graphical Model of Keystone Predators in Food Webs: Trophic Regulation of Abundance, Incidence, and Diversity Patterns in Communities. The American Naturalist 147:784–812.

Leibold, M. A. 1999. Biodiversity and nutrient enrichment in pond plankton communities. Evolutionary Ecology Research 1:73–95.

Louette, G., and L. De Meester. 2007. Predation and priority effects in experimental zooplankton communities. Oikos 116:419–426.

Mouquet, N., and M. Loreau. 2003. Community Patterns in Source-Sink Metacommunities. The American Naturalist 162:544–557.

Orrock, J. L., and R. J. Fletcher Jr. 2005. Changes in Community Size Affect the Outcome of Competition. The American Naturalist 166:107–111.

Paine, R. T. 1966. Food Web Complexity and Species Diversity. The American Naturalist 100:65–75.

Ryberg, W. A., and J. M. Chase. 2007. Predator-Dependent Species-Area Relationships. The American Naturalist 170:636–642.

Shmida, A., and M. V. Wilson. 1985. Biological Determinants of Species Diversity. Journal of Biogeography 12:1–20.

Vellend, M., D. S. Srivastava, K. M. Anderson, C. D. Brown, J. E. Jankowski, E. J. Kleynhans, N. J. B. Kraft, A. D. Letaw, A. A. M. Macdonald, J. E. Maclean, I. H. Myers-Smith, A. R. Norris, and X. Xue. 2014. Assessing the relative importance of neutral stochasticity in ecological communities. Oikos 123:1420–1430.
