## Appendix S2 for "Spatial community variability: Interactive effects of predators and isolation on stochastic community assembly"

### Appendix S2 – Analysis of expected Beta-Diversity

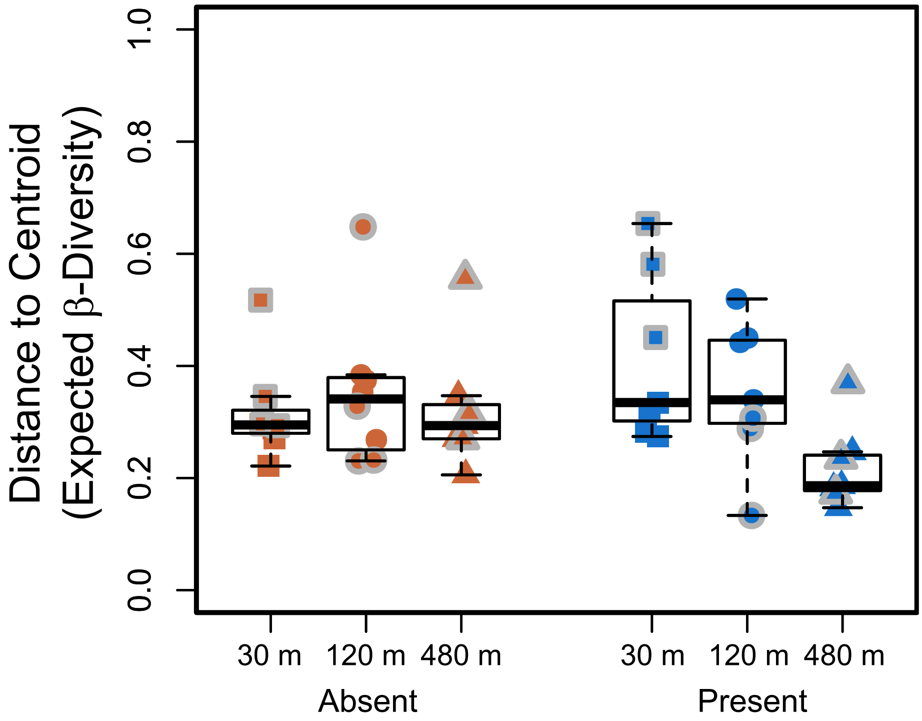


**Figure S2.1.** Box plots of values of expected distance to the centroid of 1000 randomly generated communities keeping community sizes, mean alpha diversity per treatment and number of species per treatment constant. Squares, circles, and triangles are 30 m, 120 m and 480 m isolation treatments respectively. Orange squares, circles, and triangles are fishless ponds, whereas blue are ponds with fish. Squares, circles, and triangles without a grey border are from the second survey whereas the ones with a gray border are from the third survey. Asterisks show significant differences among pairs of treatments. Because we were interested in accessing different effects of isolation in ponds with and without fish, pairwise comparisons were only done between isolation treatments within fish and fishless ponds. Pairwise comparisons between among treatments for ponds with and without fish separately showed that 30m treatments were significantly different from 480 m for ponds with fish.

**Table S2.1.** Deviance table of type II Wald Chi-square tests for the linear mixed models for expected beta-diversity. Only significant differences in Post-hoc comparisons are shown.

| | Df | Chi-square | p | Post-hoc comparisons |
|---|---|---|---|---|
| Fish | 1 | 0.011 | 0.916 | |
| **Isolation** | **2** | **7.063** | **0.029** | 30m > 480m |
| Sampling Survey | 1 | 3.639 | 0.056 | |
| **Fish : Isolation** | **2** | **8.425** | **0.015** | Fish: 30m > 480m |
| Fish : Sampling Survey | 1 | 0.170 | 0.680 | |
| **Isolation : Sampling Survey** | **2** | **12.700** | **0.002** | Second Survey: 120m > 480m<br>Third Survey: 30 >120 = 480m |
| **Fish : Isolation : Sampling Survey** | **2** | **6.840** | **0.033** | Second Survey, Fish: 120m > 480m<br>Third Survey, Fish: 30 >120 = 480m |
