## Appendix S3 for "Spatial community variability: Interactive effects of predators and isolation on stochastic community assembly"

**Appendix S3 - Results for the first sampling survey only.**

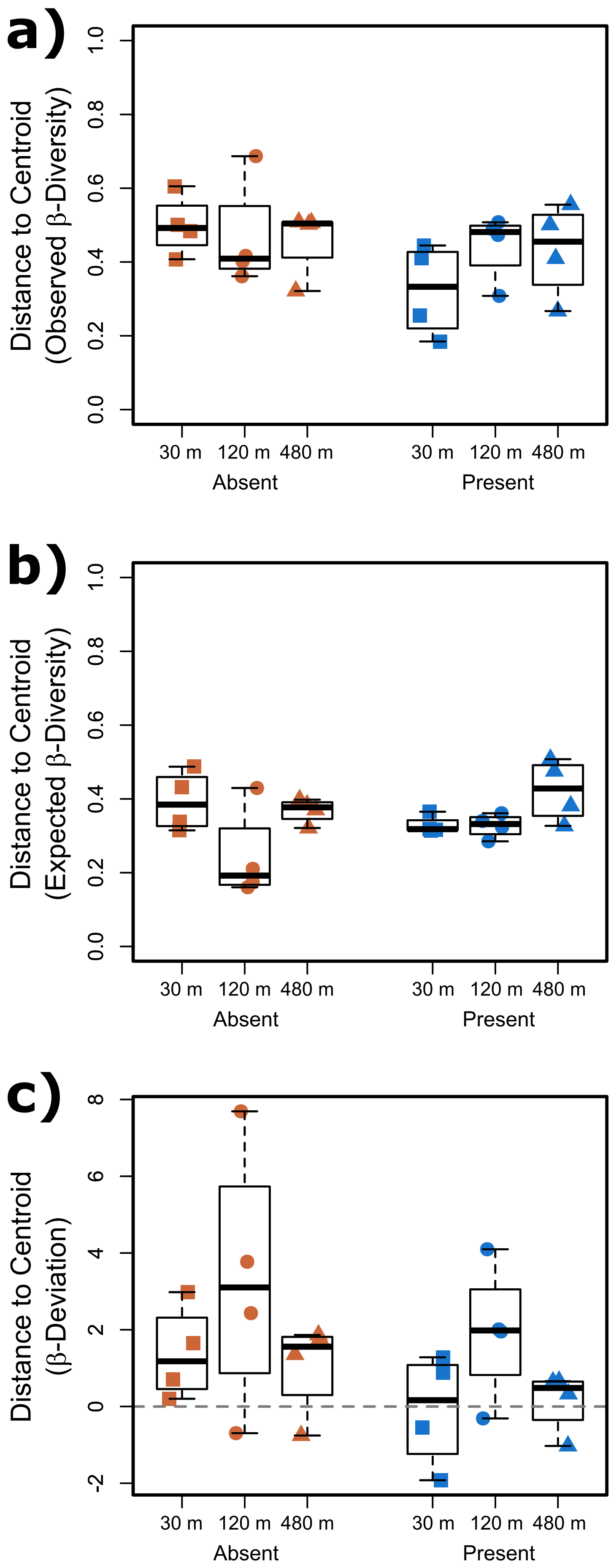


**Figure S3.1.** Box plots of values of distance to the centroid of observed dissimilarity values based on Bray-Curtis distance (a), expected distance according to 1000 simulations of null communities (b), and distance to centroid based on beta-deviation measures (c) for only the first sampling survey. Squares, circles, and triangles are 30 m, 120 m and 480 m isolation treatments respectively. Orange squares, circles, and triangles are fishless ponds, whereas blue are ponds with fish. Squares, circles, and triangles without a grey border are from the second survey whereas the ones with a gray border are from the third survey.

**Table S3.1.** Anova table of linear models for values of observed, expected beta-diversity and beta-deviation for the first sampling survey only.

| | | Observed Beta Diversity | | Expected Beta-Diversity | | Beta-Deviation | |
|---|---|---|---|---|---|---|---|
| | Df | F | p | F | p | F | p |
| Fish | 1 | 2,611 | 0.124 | 0.690 | 0.417 | 2,643 | 0.121 |
| Isolation | 2 | 0.338 | 0.717 | **4,737** | **0.022** | 3,009 | 0.075 |
| Fish : Isolation | 2 | 1,190 | 0.327 | 2,296 | 0.129 | 0.050 | 0.952 |
