## Appendix S4 for "Spatial community variability: Interactive effects of predators and isolation on stochastic community assembly"

### Appendix S4 – Local species richness

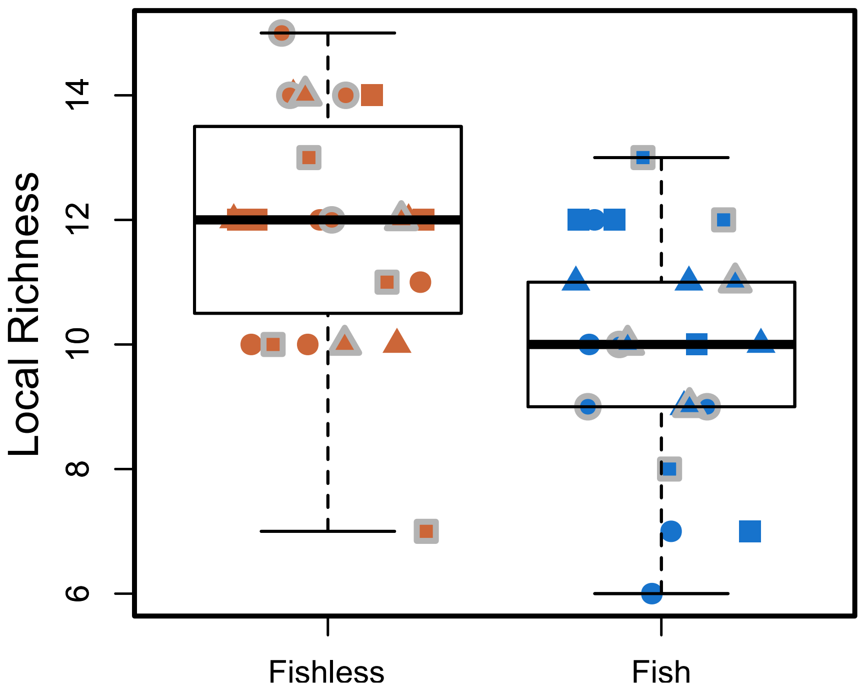


**Figure S4.1.** Box plot of local species richness in ponds with and without fish. Squares, circles, and triangles are 30 m, 120 m and 480 m isolation treatments respectively. Orange squares, circles, and triangles are fishless ponds, whereas blue are ponds with fish. Squares, circles, and triangles without a grey border are from the second survey whereas the ones with a gray border are from the third survey.
