## Appendix S5 for "Spatial community variability: Interactive effects of predators and isolation on stochastic community assembly"

### Appendix S5 - Repeating beta-diversity analyses but with a null model that does not keep gamma diversity of each treatment constant.

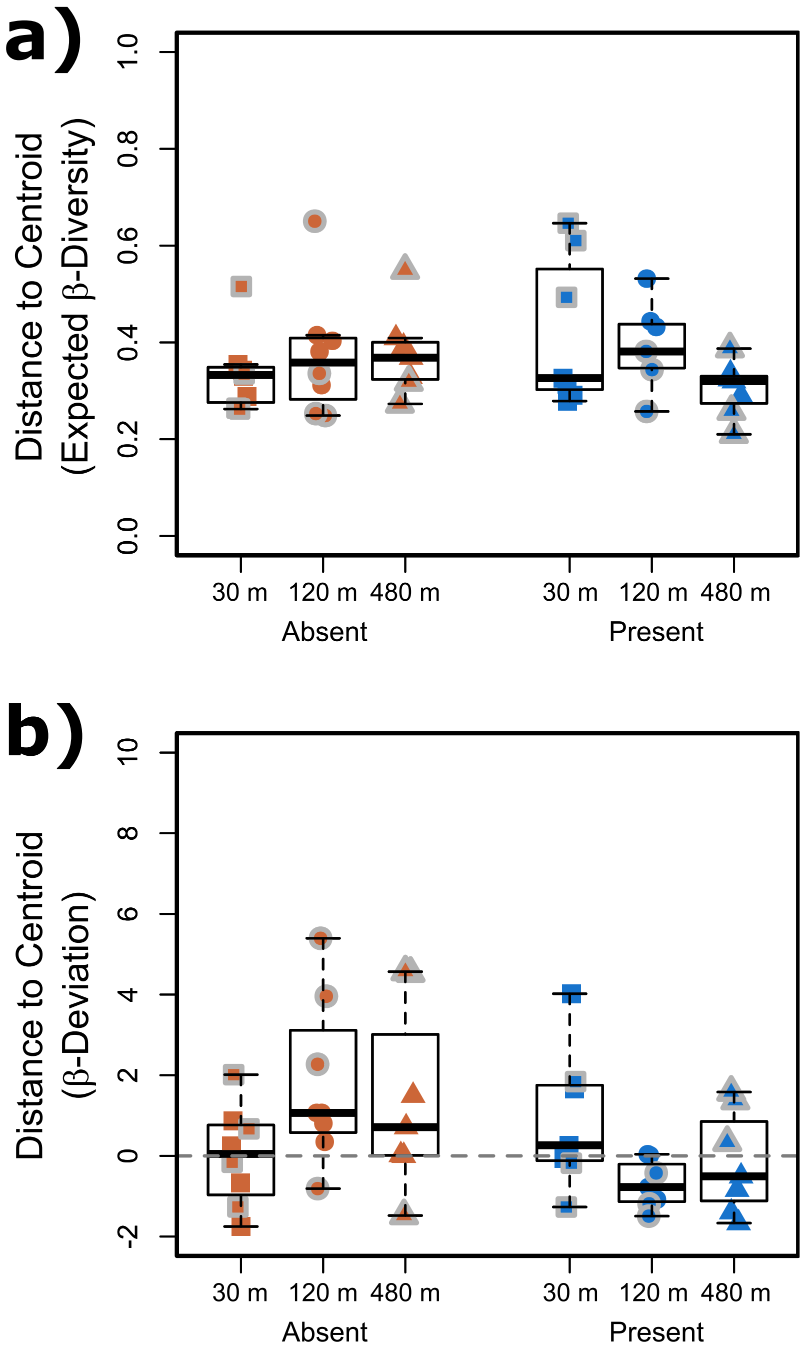

**Figure S5.1.** Box plots of values of expected beta-diversity based on 1000 randomly generated communities keeping community sizes, but not mean alpha diversity per treatment and number of species per treatment (a), and beta-deviation measures (b). Squares, circles, and triangles are 30 m, 120 m and 480 m isolation treatments respectively. Orange squares, circles, and triangles are fishless ponds, whereas blue are ponds with fish. Squares, circles, and triangles without a grey border are from the second survey whereas the ones with a gray border are from the third survey. Pairwise comparisons did not show significant differences among treatments for beta deviation.

**Table S5.1.** Deviance table of type II Wald Chi-square tests for the linear mixed models. of expected beta-diversity and beta-deviation using with a null model that does not keep gamma diversity of each treatment constant. Only significant differences in Post-hoc comparisons are shown.

| | Df | Chi-square | p | Post-hoc comparisons |
|---|---|---|---|---|
| *Expected Beta-diversity* | | | | |
| Fish | 1 | 0.222 | 0.637 | |
| Isolation | 2 | 1.895 | 0.388 | |
| Sampling Survey | 1 | 0.708 | 0.400 | |
| **Fish : Isolation** | **2** | **6.263** | **0.044** | Fish: 30m > 480m |
| Fish : Sampling Survey | 1 | 0.614 | 0.433 | |
| **Isolation : Sampling Survey** | **2** | **9.222** | **0.010** | Third: 30m > 480m |
| **Fish : Isolation : Sampling Survey** | **2** | **8.756** | **0.013** | Third, Fish: 30m > 120m = 480m |
| *Beta-deviation* | | | | |
| **Fish** | **1** | **4.215** | **0.040** | |
| Isolation | 2 | 0.247 | 0.884 | |
| Sampling Survey | 1 | 3.392 | 0.066 | |
| **Fish : Isolation** | **2** | **8.105** | **0.017** | |
| Fish : Sampling Survey | 1 | 2.620 | 0.106 | |
| Isolation : Sampling Survey | 2 | 4.941 | 0.085 | |
| Fish : Isolation : Sampling Survey | 2 | 1.813 | 0.404 | |
