## Appendix S6 for "Spatial community variability: Interactive effects of predators and isolation on stochastic community assembly"

### Appendix S6 - Results for predators and non-predators separately.

**Table S6.1.** Deviance table of type II Wald Chi-square tests for generalized linear mixed models of abundance and local richness for predatory insects only. Only significant differences in Post-hoc comparisons are shown.

| | Df | Chi-square | p | Post-hoc comparisons |
|---|---|---|---|---|
| *Abundance* | | | | |
| **Fish** | **1** | **62.925** | **<0.001** | |
| **Isolation** | **2** | **50.169** | **<0.001** | 30m > 120m = 480m |
| **Sampling Survey** | **1** | **9.412** | **0.002** | |
| Fish : Isolation | 2 | 4.946 | 0.084 | |
| **Fish : Sampling Survey** | **1** | **6.294** | **0.012** | Second: Fishless > Fish<br>Third: Fishless >> Fish |
| Isolation : Sampling Survey | 2 | 5.145 | 0.076 | |
| Fish : Isolation : Sampling Survey | 2 | 4.124 | 0.127 | |
| *Local Richness* | | | | |
| **Fish** | **1** | **20.258** | **<0.001** | |
| **Isolation** | **2** | **13.628** | **0.001** | 30m > 120m = 480m |
| Sampling Survey | 1 | 0.601 | 0.438 | |
| Fish : Isolation | 2 | 3.361 | 0.186 | |
| Fish : Sampling Survey | 1 | 1.415 | 0.234 | |
| Isolation : Sampling Survey | 2 | 3.296 | 0.192 | |
| **Fish : Isolation : Sampling Survey** | **2** | **11.199** | **0.004** | Second, Fish: 30m > 120m<br>Third, Fish: 30m > 120m = 480m |

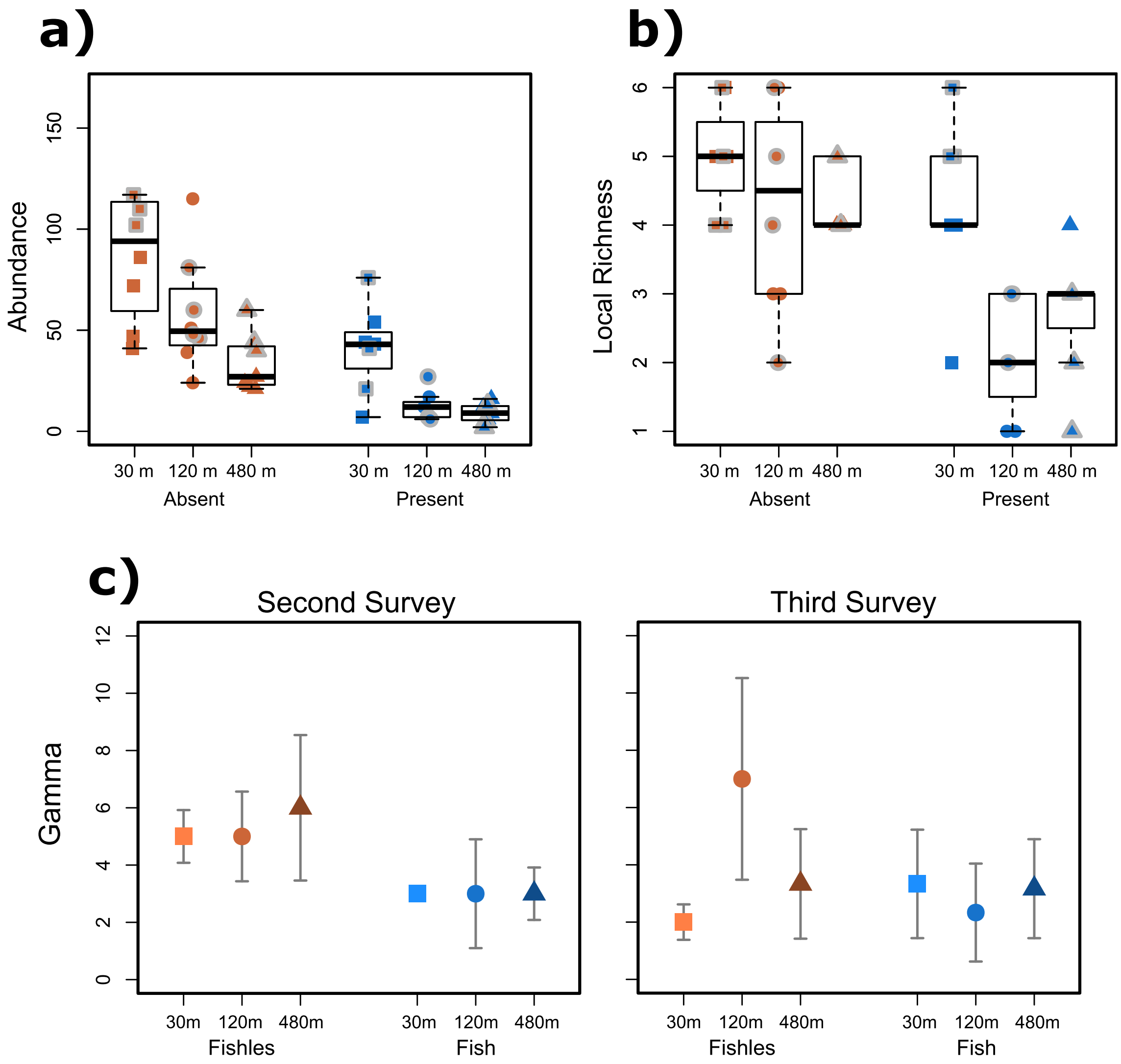

**Figure S6.1.** Box plot of abundance (i.e. community size) (a) and local richness (b) for predatory insects only. Squares, circles, and triangles are 30 m, 120 m and 480 m isolation treatments respectively. Orange squares, circles, and triangles are fishless ponds, whereas blue are ponds with fish. Squares, circles, and triangles without a grey border are from the second survey whereas the ones with a gray border are from the third survey. (c) Gamma diversity (regional richness) per treatment in the second (left panel) and third surveys (right panel) considering four sample units per treatment. Grey bars are estimated 95% confidence intervals. Estimates whose confidence intervals do not overlap are significantly different.

**Table S6.2.** Deviance table of type II Wald Chi-square tests for generalized linear mixed models of abundance and local richness for non-predatory insects only. Only significant differences in Post-hoc comparisons are shown.

| | Df | Chi-square | p | Post-hoc comparisons |
|---|---|---|---|---|
| *Abundance* | | | | |
| Fish | 1 | 0.823 | 0.364 | |
| Isolation | 2 | 1.831 | 0.400 | |
| **Sampling Survey** | **1** | **22.131** | **<0.001** | |
| Fish : Isolation | 2 | 2.242 | 0.326 | |
| Fish : Sampling Survey | 1 | 2.524 | 0.112 | |
| **Isolation : Sampling Survey** | **2** | **7.413** | **0.025** | 120m and 480m: Second < Third |
| Fish : Isolation : Sampling Survey | 2 | 1.218 | 0.544 | |
| *Local Richness* | | | | |
| Fish | 1 | 0.975 | 0.323 | |
| Isolation | 2 | 5.542 | 0.063 | |
| Sampling Survey | 1 | 0.044 | 0.834 | |
| Fish : Isolation | 2 | 0.757 | 0.685 | |
| Fish : Sampling Survey | 1 | 0.238 | 0.625 | |
| **Isolation : Sampling Survey** | **2** | **6.574** | **0.037** | Third: 30m > 120m |
| Fish : Isolation : Sampling Survey | 2 | 4.893 | 0.087 | |

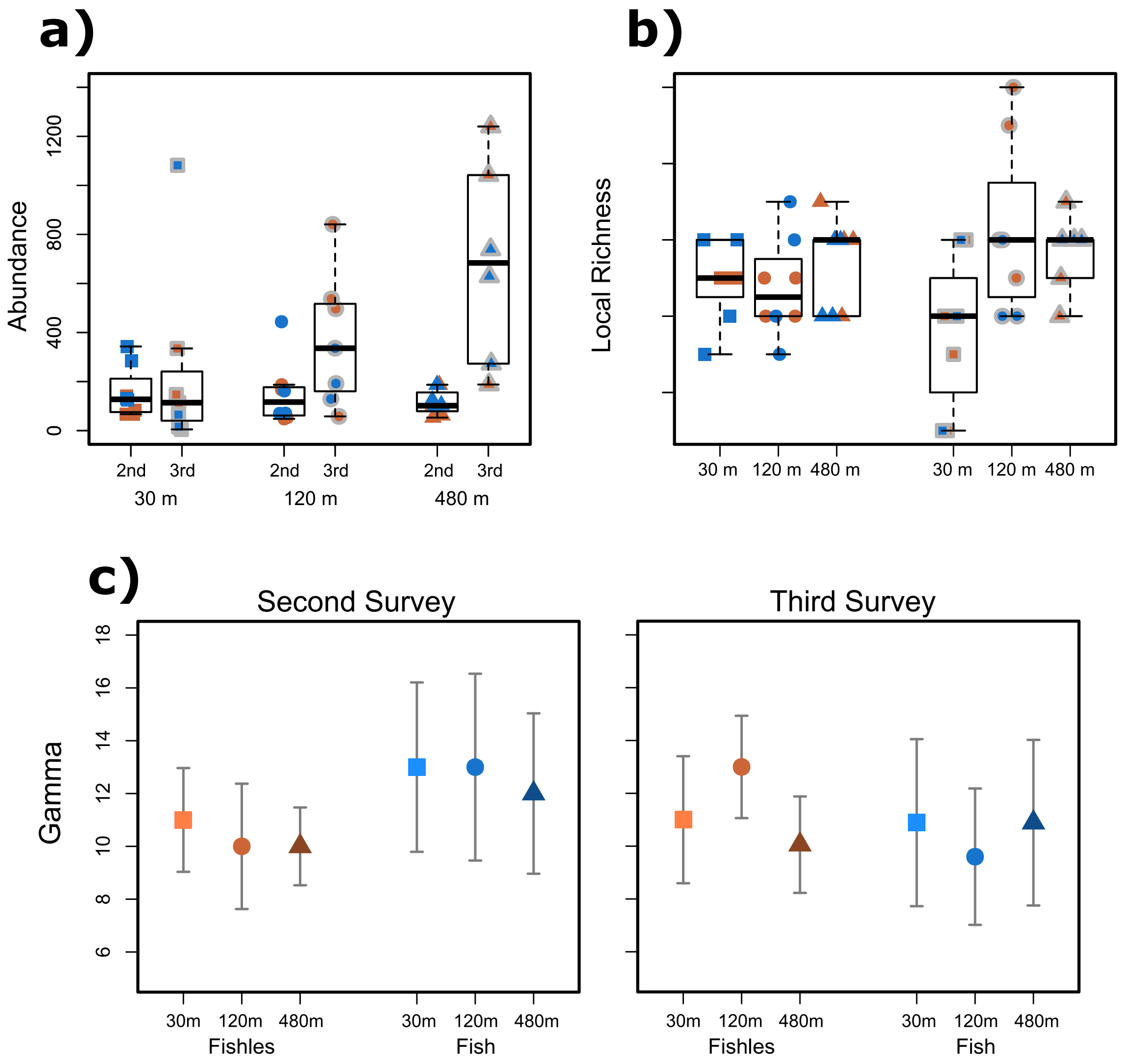

**Figure S6.2.** Box plot of abundance (i.e. community size) (a) and local richness (b) for non-predatory insects only. Squares, circles, and triangles are 30 m, 120 m and 480 m isolation treatments respectively. Orange squares, circles, and triangles are fishless ponds, whereas blue are ponds with fish. Squares, circles, and triangles without a grey border are from the second survey whereas the ones with a gray border are from the third survey. (c) Gamma diversity (regional richness) per treatment in the second (left panel) and third surveys (right panel) considering four sample units per treatment. Grey bars are estimated 95% confidence intervals. Estimates whose confidence intervals do not overlap are significantly different.

**Table S6.3.** Deviance table of type II Wald Chi-square tests of linear mixed models for observed, expected beta-diversity and beta-deviations considering only predatory insects. Only significant differences in Post-hoc comparisons are shown.

| | Df | Chi-square | p | Post-hoc comparisons |
|---|---|---|---|---|
| *Observed Beta-diversity* | | | | |
| Fish | 1 | 1.199 | 0.273 | |
| Isolation | 2 | 2.129 | 0.345 | |
| **Sampling Survey** | **1** | **5.533** | **0.019** | |
| **Fish : Isolation** | **2** | **6.370** | **0.041** | |
| **Fish : Sampling Survey** | **1** | **9.009** | **0.003** | Third: Fishless < Fish |
| **Isolation : Sampling Survey** | **2** | **7.528** | **0.023** | Third: 30m < 480m |
| Fish : Isolation : Sampling Survey | 2 | 3.697 | 0.157 | |
| *Expected Beta-diversity* | | | | |
| **Fish** | **1** | **12.353** | **<0.001** | |
| Isolation | 2 | 3.958 | 0.138 | |
| Sampling Survey | 1 | 1.930 | 0.165 | |
| Fish : Isolation | 2 | 3.541 | 0.170 | |
| **Fish : Sampling Survey** | **1** | **3.975** | **0.046** | Third: Fishless < Fish |
| **Isolation : Sampling Survey** | **2** | **6.463** | **0.039** | Third: 30m < 480m |
| **Fish : Isolation : Sampling Survey** | **2** | **6.679** | **0.035** | Third, Fish: 30 < 480m |
| *Beta-deviation* | | | | |
| **Fish** | **1** | **4.384** | **0.036** | |
| Isolation | 2 | 3.746 | 0.154 | |
| Sampling Survey | 1 | 0.912 | 0.340 | |
| Fish : Isolation | 2 | 2.067 | 0.356 | |
| **Fish : Sampling Survey** | **1** | **9.372** | **0.002** | |
| Isolation : Sampling Survey | 2 | 0.703 | 0.704 | |
| Fish : Isolation : Sampling Survey | 2 | 4.893 | 0.087 | |

**Table S6.4.** Deviance table of type II Wald Chi-square tests of linear mixed models for observed, expected beta-diversity and beta-deviations considering only non-predatory insects. Only significant differences in Post-hoc comparisons are shown.

| | Df | Chi-square | p | Post-hoc comparisons |
|---|---|---|---|---|
| *Observed Beta-diversity* | | | | |
| **Fish** | **1** | **3.887** | **0.049** | |
| Isolation | 2 | 3.178 | 0.204 | |
| Sampling Survey | 1 | 3.265 | 0.071 | |
| **Fish : Isolation** | **2** | **13.724** | **0.001** | Fish: 30m > 120m = 480m |
| Fish : Sampling Survey | 1 | 0.292 | 0.589 | |
| **Isolation : Sampling Survey** | **2** | **11.718** | **0.003** | Third: 30m > 120m = 480m |
| Fish : Isolation : Sampling Survey | 2 | 5.014 | 0.082 | |
| *Expected Beta-diversity* | | | | |
| Fish | 1 | 0.821 | 0.365 | |
| **Isolation** | **2** | **10.389** | **0.006** | 30m > 480m |
| **Sampling Survey** | **1** | **4.924** | **0.026** | |
| Fish : Isolation | 2 | 4.296 | 0.117 | |
| Fish : Sampling Survey | 1 | 0.874 | 0.350 | |
| **Isolation : Sampling Survey** | **2** | **15.799** | **<0.001** | Third: 30m > 120m = 480m |
| Fish : Isolation : Sampling Survey | 2 | 1.834 | 0.400 | |
| *Beta-deviation* | | | | |
| **Fish** | **1** | **4.977** | **0.026** | |
| **Isolation** | **2** | **6.650** | **0.036** | |
| Sampling Survey | 1 | 0.074 | 0.785 | |
| **Fish : Isolation** | **2** | **7.499** | **0.024** | Fishless: 30m < 120m = 480m |
| Fish : Sampling Survey | 1 | 0.197 | 0.657 | |
| Isolation : Sampling Survey | 2 | 3.644 | 0.162 | |
| **Fish : Isolation : Sampling Survey** | **2** | **13.242** | **0.001** | Second, Fishless: 30m = 120m < 480m<br>Third, Fishless: 30m < 120m |

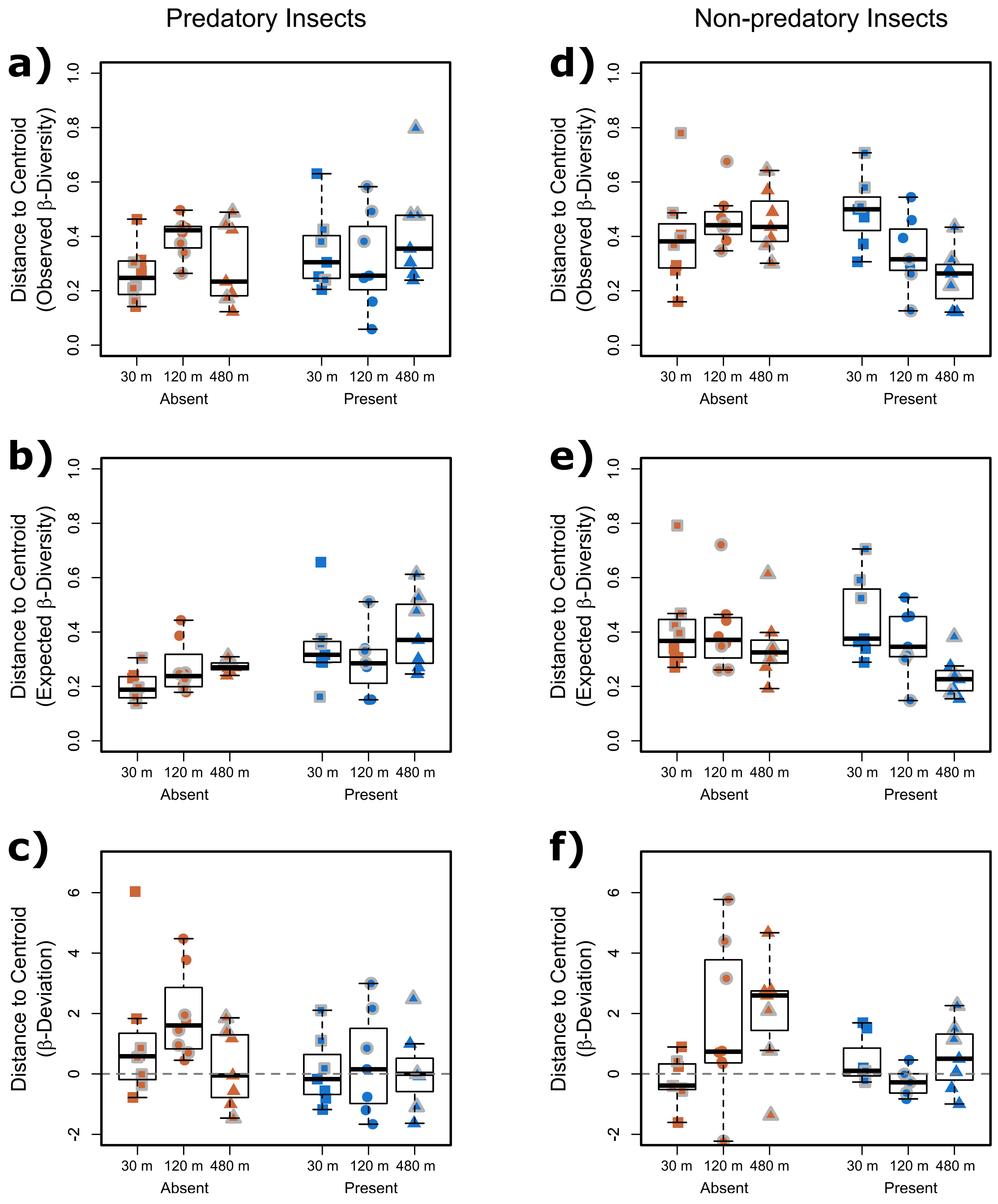

**Figure S6.3.** Box plots of values of distance to the centroid of observed dissimilarity values based on Bray-Curtis distance (a), expected distance according to 1000 simulations of null communities (b), and distance to centroid based on beta-deviation measures (c) for predatory insects only (a, b and c) and consumers only (d, e and f). Squares, circles, and triangles are 30 m, 120 m and 480 m isolation treatments respectively. Orange squares, circles, and triangles are fishless ponds, whereas blue are ponds with fish. Squares, circles, and triangles without a grey border are from the second survey whereas the ones with a gray border are from the third survey. Pairwise comparisons of observed beta-diversity and beta-deviation for predatory showed that both were higher in the last survey, but only for ponds with fish. For consumers, pairwise comparisons of observed beta-diversity showed that 30 m ponds were different from 480 m ponds, but only for ponds with fish. For beta-deviation pairwise comparisons showed no significant difference among treatments.
